# Myocardial damage induced by a single high dose of isoproterenol in C57BL/6J mice triggers a persistent adaptive immune response against the heart

**DOI:** 10.1101/2020.02.27.962696

**Authors:** Elvira Forte, Mona Panahi, Fu Siong Ng, Joseph J. Boyle, Jane Branca, Olivia Bedard, Muneer G. Hasham, Lindsay Benson, Sian E. Harding, Nadia Rosenthal, Susanne Sattler

## Abstract

Heart failure is the common final pathway of a range of conditions resulting in myocardial damage and a major cause of morbidity and mortality worldwide. Strategies to improve tissue repair and prevent heart failure thus remain an urgent clinical need. Recent studies have documented activation of the adaptive immune system in response to myocardial damage and have implicated anti-heart autoimmunity in the development of heart failure. In an attempt to target anti-heart autoimmune responses as new therapeutic avenue, the number of experimental studies using *in vivo* models of myocardial damage to study the ensuing immune response has surged.

The beta-adrenergic agonist isoproterenol-hydrochloride has been used for its cardiac effects in a variety of different dosing and administration regimes. Most prominently, low doses (<10mg/kg sc) over an extended time period induce cardiac hypertrophy and fibrosis. In addition, single injections of high doses (>100mg/kg) induce cardiomyocyte necrosis and have been used to mimic acute myocardial necrotic lesions as seen in myocardial infarction (MI). However, despite significant resource and animal welfare advantages, concerns about off-target effects and clinical relevance have so far limited uptake in the cardiovascular research community.

To assess suitability of the isoproterenol model for the analysis of chronic post-MI immunological readouts, we treated C57BL/6J mice with a single intra-peritoneal bolus injection of 160mg/kg isoproterenol. Our results confirm the presence of necrotic lesions in the myocardium with significant resemblance of the histopathology of Type 2 MI. Kidneys develop mild fibrosis secondary to early cardiac damage, while other organs remain unaffected. Most importantly, we showed that isoproterenol treatment causes myocardial inflammation and fibrosis, activation of T cells in the heart-draining mediastinal lymph nodes, deposition of mature antibodies in the myocardium and the presence of auto-antibodies against the heart in the serum 12 weeks after the initial injury.

In summary, this simple and cost-effective experimental model with significant animal welfare benefits induces myocardial damage reminiscent of human Type 2 MI, which is followed by a persistent adaptive immune response against the heart. This makes it a suitable and high-throughput model to study pathological mechanisms of anti-heart autoimmunity as well as to test potential immunomodulatory therapeutic approaches.

## Introduction

Clinical guidelines distinguish between myocardial injury and myocardial infarction (MI). While myocardial injury encompasses any type of acute myocardial damage such as necrosis induced by myocarditis, sepsis or endogenous catecholamines, MI is defined as ischemic damage resulting from an insufficient myocardial oxygen supply ^1 2^. Aherosclerotic vascular disease is a prominent cause of MI, where disruption of an atherosclerotic plaque can prevent coronary blood flow through a given artery resulting in infarction of the dependent myocardium. This is referred to as Type 1 MI (T1MI). Type 2 MI (T2MI) is a heterogeneous syndrome that occurs in the absence of coronary artery disease (CAD) and an acute atherothrombotic coronary event. Instead, other conditions such as hypoxemia, hypo/hypertension, tachycardia or tachyarrhythmias lead to an imbalance between oxygen supply and demand ^3 4^. T2MI is a common but under-recognised clinical entity, with prevalence estimates of up to 58% of MI patients ^5^.

Importantly, improved intervention and treatment after myocardial damage have significantly increased immediate survival. However, the regenerative capacity of the adult heart is minimal and healing is achieved by fibrotic repair which replaces damaged myocardium with non-contractile scar tissue. This impairs cardiac function and 62.7% of MI patients develop heart failure within 6 years after infarction ^6^. The early immune response to myocardial necrosis is crucial for quick tissue repair, but excessive inflammation is involved in the development of heart failure ^7 8 9 10^. Large amounts of cardiac self-antigens, including myosin and troponin, in a highly inflammatory environment activate the adaptive immune system. Auto-reactive T and B cells can be long-lived and have been suggested to cause ongoing low level tissue destruction ^11 12^ hampering regenerative efforts and exacerbating development towards heart failure ^13 14^. There is an urgent need for better recognition and understanding of these autoimmune processes to prevent further myocardial damage.

Investigation of the adaptive immune response to cardiac damage is a highly complex multi-dimensional process, which to date still needs *in vivo* experimentation. In rodents, surgical coronary artery ligation most prominently of the left anterior descending artery (LAD), is the most commonly used experimental model for human T1MI ^15^. LAD ligation is induced during open chest surgery, an invasive low-throughput procedure associated with high mortality rates. The use of the ß-adrenergic agonist isoproterenol to induce cardiotoxicity and infarct-like lesions has been described over 50 years ago in rats ^16 17^. Fibrosis in response to isoproterenol treatment is related to cardiomyocyte necrosis. Necrotic cardiomyocytes are mostly localised in the sub-endocardium, numbers peak at 24h and debris is removed by 48h ^18^. Isoproterenol is now routinely used as a long term low-dose treatment to induce cardiac hypertrophy ^19 20 21^, and few research groups have also used it to induce acute infarct-like lesions by a one-off injection of higher doses ^22 23 24^. Importantly, the isoproterenol model recapitulates several of the salient feature of T2MI. Positive inotropic and chronotropic effects lead to increased myocardial oxygen consumption consumption and Ca^2+^ leakage causing cardiomyocyte necrosis ^24 25^, while tachycardia leads to a reduction in diastolic perfusion time and coronary blood flow ^26^, collectively resulting in patchy acute myocardial damage and necrosis. Notably, it also mechanistically and pathologically models a pattern of injury termed inotrope-induced cardiac damage or pressor-injury or catecholamine-heart, seen in the cardiac transplant and terminal heart failure setting ^27^.

Here we characterise the immunological effects of a one-off intra-peritoneal injection of isoproterenol as a simple, resource-efficient, highly reproducible and significantly less invasive alternative to surgical MI induction. We show that T cell activation in the mediastinal lymph nodes and anti-heart auto-antibody production can be investigated without the need for invasive surgical procedures, as the adaptive immune system will respond to cardiac antigens released from necrotic tissue, irrespective of the original trigger of necrosis.

## Materials and Methods

### Mice

All animal procedures carried out at Imperial College London were approved by the Imperial College Governance Board for Animal Research and in accordance with the UK Home Office Animals (Scientific Procedures) Act 1986 and Directive 2010/63/EU of the European Parliament on the protection of animals used for scientific purposes. All animal work carried out at The Jackson Laboratories were approved by The Jackson Laboratory Institutional Animal Care and Use Committee and were in accordance with national and international regulations. Mice used were 8-to 12-week-old C57BL/6J males and females; control mice were age- and sex-matched. Mice were housed under SPF conditions (Imperial College London) or in conventional cages (The Jackson Laboratories) in temperature-controlled facilities on a 12h light/dark cycle on standard diet.

### Isoproterenol treatment and tissue harvest

Mice were treated with isoproterenol HCL (Sigma-Aldrich, St. Louis, MO, USA) in Dulbecco’s phosphate-buffered-saline (DPBS; Sigma-Aldrich) by intraperitoneal injection once at a dose of 160 mg/kg. Control mice were treated with DPBS at equivalent volume. Tissue was harvested 1 or 2 weeks post-injection. On the day of tissue collection, 200 ul blood was collected. Blood was incubated on ice for 30 minutes, then centrifuged 3 minutes at max rcf. Serum was collected and stored at −20 ºC for later use. Hearts and lymph nodes were isolated after *in situ* perfusion with ice-cold DPBS supplemented with 0.9 mM CaCl2 through the apex of the left ventricle of the heart to clear blood from heart chambers and blood vessels.

### Histology and scoring of damage parameters

Organs of treated and untreated mice were excised after perfusion as described above, fixed in 10% neutral buffered formalin overnight, and stored in 70% ethanol. For wax-embedding and histology, tissue samples were dehydrated in an increasing gradient of ethanol and embedded in paraffin. Five μm sections were cut and de-waxed and rehydrated in an ethanol gradient. Sections were stained with hematoxylin and eosin (H&E) and Picrosirius Red. All reagents were purchased from Sigma Aldrich (Sigma-Aldrich, Dorset, UK). Semi-quantitative scoring of heart sections was performed as established previously ^28^. Hematoxylin & eosin stained sections were used to analyse and score mononuclear cell infiltration in hearts. Picrosirius Red staining was used to analyse and score fibrosis in hearts and kidneys. Individual parameters were scored on a scale of 0 (none), 1 (mild), 2 (moderate) to 3 (severe). Scores were obtained from 4 areas each on two midline cross sections per animal by a blinded researcher. Images were captured using a LMD7000 microscope (Leica microsystems, Milton Keynes, UK) and processed for quantification of nuclei (cell count) and area of fibrosis using the public domain software ImageJ (NIH; http://rsb.info.nih.gov,)^29^.

### cTroponin and anti-heart autoantibody ELISA

A mouse cardiac troponin I (cTI) ELISA Kit (MyBioSource, San Diego, CA, USA) was used to determine cTI concentrations in post-ISOPROTERENOL serum as instructed by the manufacturer. Serum samples were diluted 2-fold in supplied diluent. A standard curve was generated to calculate cTI concentrations in pg/ml.

The ELISA protocol for detection of **mouse anti-heart auto-antibodies** was optimised as described previously ^28^. ELISA plates (SpectraMax Paradigm Molecular-Devices, UK) were coated with 50μl per well of 4μg/μl pig heart lysate (Novus Biologicals, Bio-Techne, Abingdon, UK) diluted in PBS overnight at 4 ºC. Plates were washed three times for 5 minutes each with 200 μl per well of ELISA washing buffer. 50 μl of post-ISOPROTERENOL serum was added diluted 1:10 and 1:100 for overnight incubation at 4 ºC. Detection reagents were anti-mouse IgG-HRP (all BioLegend, London, UK). Optical densities were measured at 450 nm using a SpectraMAX i3 microplate reader (Molecular Devices, San Jose, CA, USA).

### Mediastinal lymph node flow cytometry

For isolation of single cells, mediastinal lymph nodes were manually minced through 70 μm filters using the rubber end of a 1ml syringe. Red blood cells were lysed using Red Blood Cell Lysis buffer (Sigma-Aldrich, Dorset, UK) according to the manufacturer’s instructions. The obtained cell mixture was then used for flow cytometric analysis. Antibodies used: anti-mouse CD45 (CD45-APC-Cy7 - cat. 103116, clone 30-F11, LOT B185138, dilution 1:800), anti-mouse CD3 (CD3-APC - cat. 100235, clone 17A2, LOT B166471, dilution 1:200), anti-mouse CD4 (CD4-FITC - cat. 115505, clone 6D5, LOT B131781, dilution 1:200), anti-mouse CD8a (CD8a-PE - cat. 101207, clone M1/70, LOT B166034, dilution 1:400), and anti-mouse CD62L (CD62L-APC - cat. 137607, clone 29A1.4, LOT B152186, dilution 1:200) and anti-mouse CD44(CD44-APC - cat. 137607, clone 29A1.4, LOT B152186, dilution 1:200) were purchased from BioLegend (BioLegend, London, UK). Antibody dilutions in cell staining buffer containing 1 % TruStain fcX™ (anti-mouse CD16/32) Antibody (both BioLegend, London, UK) were used to stain according to the manufacturer’s protocol. Antibody validation data is available from http://www.biolegend.com. Samples were acquired using a BD LSRII (Beckton Dickinson, Oxford, UK) and analysed using Flow Jo version 10.6.1 (Treestar, Ashland, OR, USA) software.

### Immunohistochemistry

For detection of in vivo antibody deposition in cardiac tissue, 5 μm sections of frozen post-ISOPROTERENOL hearts were stained with goat anti-mouse IgG-FITC (cat. F5387, Sigma-Aldrich, Dorset, UK), rat anti-mouse IgM-PE (cat. 406507, BioLegend, London, UK). Images were captured using a LMD7000 microscope (Leica microsystems, Milton Keynes, UK) processed using the public domain software ImageJ (NIH; http://rsb.info.nih.gov) (Schenider et al 2012)

#### Statistical analysis

Statistical analysis was performed using GraphPad Prism 8 and data were presented as mean±s.e.m throughout. Comparison between 2 groups was performed using student’s t-test. Comparison between multiple experimental groups was performed using one- or two-way ANOVA with Dunnett's multiple comparisons post hoc test to obtain multiplicity-adjusted p-values. Non-parametric scoring data were analysed using Kruskal-Wallis test with Dunn's multiple comparisons post hoc test to obtain multiplicity-adjusted p-values. Differences were considered significant at p<0.05.

## Results

### (1) A single dose of 160mg/kg isoproterenol induces significant cardiac damage in C57BL/6J mice

Different mouse strains show variable susceptibility to induction of cardiac damage and fibrosis, and C57BL/6J mice have a very robust cardiac phenotype and are resistant to myocardial damage and cardiac fibrosis ^30 31^. They are among the best characterised mouse strains and commonly used for genetic modification ^32^. We therefore performed a dose titration experiment to investigate if C57BL/6J mice were susceptible to developing cardiac lesions while maintaining high animal welfare standards. Mice were treated with a one-off bolus intra-peritoneal injection of increasing doses of isoproterenol starting with 40mg/kg up to 320mg/kg. The highest dose of 320mg/kg was administered as 2 separate 160mg/kg doses on subsequent days. Isoproterenol effects were evident from 15mins after injection, at which point mice ceased moving and increased respiration rates were observed. In general, mice recovered within 2 hours as judged by the return of normal feeding and grooming behaviour. 160mg/kg isoproterenol induced significant increase in the cardiac damage biomarker cardiac Troponin I in the serum (Figure 1A) as well as myocardial infiltration and fibrosis 2 weeks after challenge (Figure 1B). Spleen/body ratio as a measure of systemic inflammation and body weight as indicator of overall health was not affected with doses up to 160mg/kg (Figure 1C, D). 160mg/kg each on 2 subsequent days however increased spleen/body ratio and reduced body weight. A single injection of 160mg/kg was thus chosen as standard dose for subsequent experiments. Histopathologically, cardiac damage was accompanied by early patterns of necrosis followed by a maturing repair response resulting in granulation tissue and eventually fibrosis over an appropriate time-span of weeks. Consistent with catecholaminergic damage, and a strong component of ischemia, injury was most prominent in the subendocardium of the left ventricular free wall, but was essentially confluent and transmural in the septum (Figure 1E).

**Figure 1:**
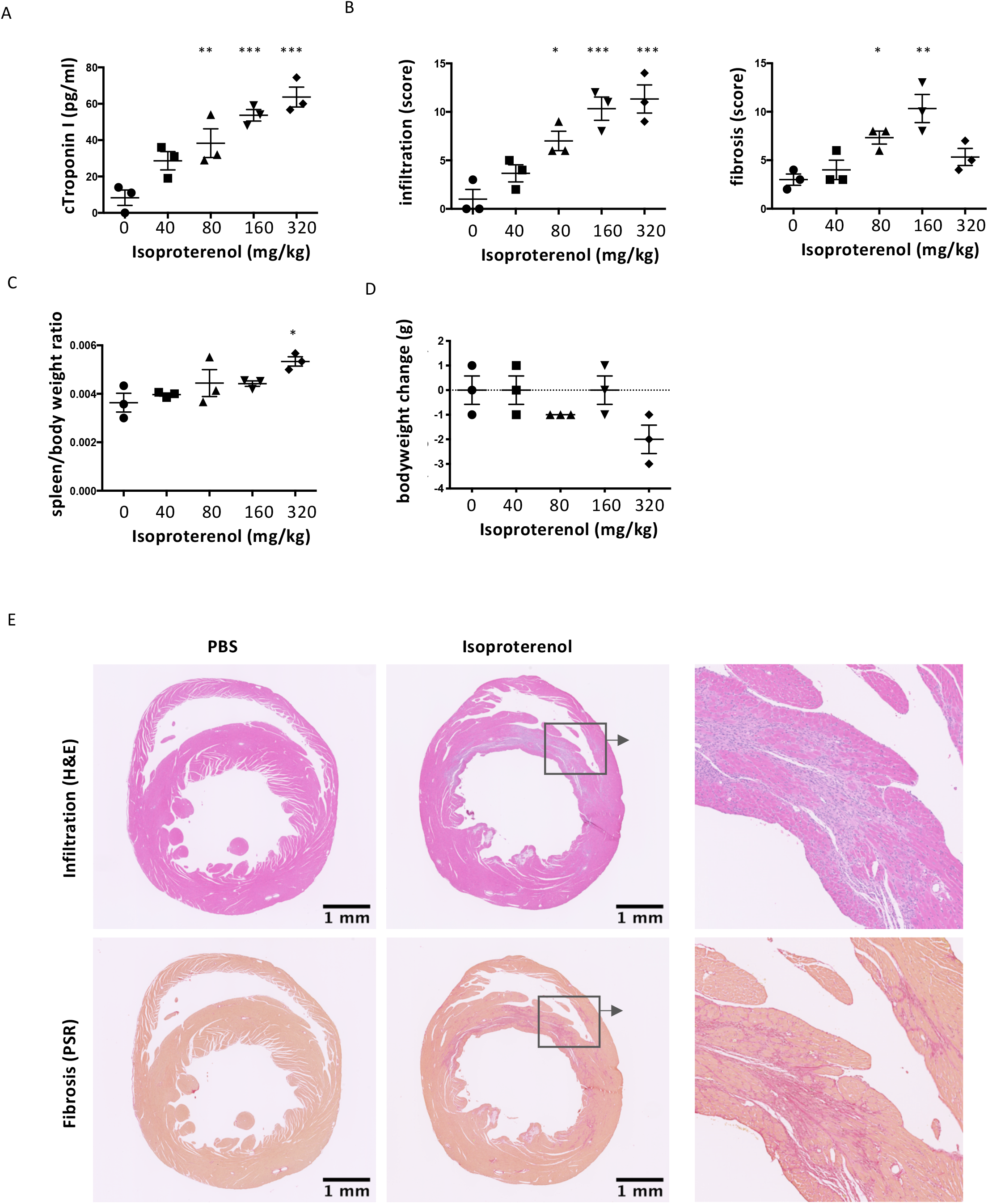
A one-off high dose of isoproterenol induces myocardial inflammation and fibrosis in C57Bl/6JJ mice. C57BL/6J mice were treated with a single increasing one-off dose of isoproterenol to induce myocardial ischemia and cardiomyocyte necrosis and analysed 2 weeks after challenge. (A) Serum cardiac troponin I levels to confirm myocardial injury. (B) Infiltration and fibrosis were scored on a scale from 0 to 3 (none, mild, moderate, severe) in 4 fields of view in 2 heart cross-sections at mid-level per mouse. (C) Splenomegaly (spleen/body weight ratio) as a measure of systemic immune activation. (D) Change in body weight. (E) Micrographs of H&E- and Picrosirius Red-stained paraffin-embedded cross sections of the hearts showing mononuclear cell infiltration (H&E) and interstitial fibrosis (Picrosirius Red). n=3/group. Data are expressed as mean +/− s.e.m., *p<0.05, **p<0.001, ***p<0.0001 (B: non-parametric Kruskal-Wallis with Dunn's multiple comparisons post hoc test comparing each time point to baseline (A,C,D) one-way ANOVA with Dunnett's multiple comparisons post hoc test comparing each time point to baseline).

To further improve animal welfare benefits of the isoproterenol model, we included pre-treatment with 0.05mg/kg buprenorphine for pain relief. Buprenorphine was injected subcutaneously 30 minutes before isoproterenol administration. Manual mouse grimace scale (MGS) scoring was deemed not suitable, as no orbital tightening, or changes in nose and cheek bulge and whisker position was notable ^33^. Instead, we measured activity (moving, feeding, drinking, grooming) as a reliable sign of both the onset of and the recovery from acute isoproterenol effects (Figure 2A). Adding buprenorphine to the isoproterenol treatment shortened the average time to recovery by 20 minutes without affecting the degree of cardiac inflammatory and fibrotic damage (Figure 2B).

**Figure 2:**
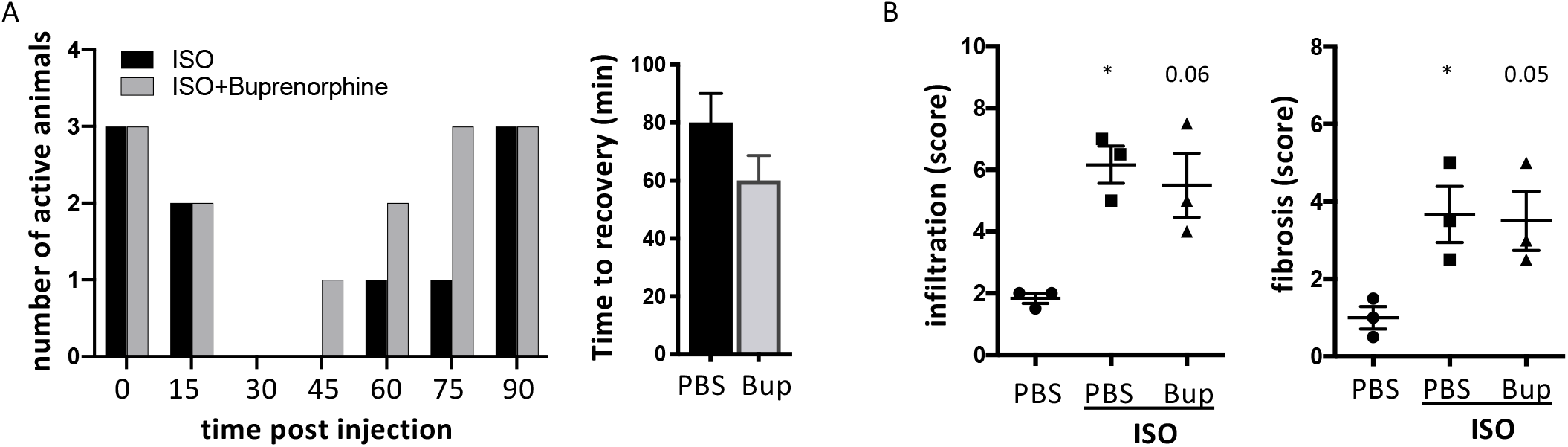
Addition of 0.05mg/kg buprenorphine further decreases procedural severity as measured by activity scoring without affecting infiltration and fibrosis scores. C57BL/6J mice were treated with a single 160mg/kg dose of isoproterenol with and without 0.05mg/kg buprenorphine to induce myocardial ischemia and cardiomyocyte necrosis under analgesia. (A) Count of animals with normal movement and activity. (B) Average time until recovery of normal activity pattern. (C) Infiltration and fibrosis in cardiac tissue sections were scored on a scale from 0 to 3 (none, mild, moderate, severe) in 4 fields of view in 2 heart cross-sections at mid-level per mouse. n=3/group. Histopathology scoring data are expressed as mean +/− s.e.m., *p<0.05 (non-parametric Kruskal-Wallis with Dunn's multiple comparisons post hoc test comparing each time point to baseline).

### (2) Kidney, liver, lung and skeletal muscle are protected from direct isoproterenol-induced damage

To define the degree of off-target damage to other organs, we performed a thorough histopathological assessment of kidneys, liver, lung and skeletal muscle at week 1 and 2 after injection of 160mg/kg isoproterenol. Myocardial mononuclear infiltration and fibrosis was confirmed (Figure 3A). No pathological changes were observed in any of the other tested organs in any of the mice (n=5) at week 1 (Figure 3B, C). At week 2, mild acute tubular injury and fibrosis was detectable in the kidneys (Figure 3B, C) which is likely secondary to cardiac damage as previously observed after surgical LAD ligation ^34^.

**Figure 3:**
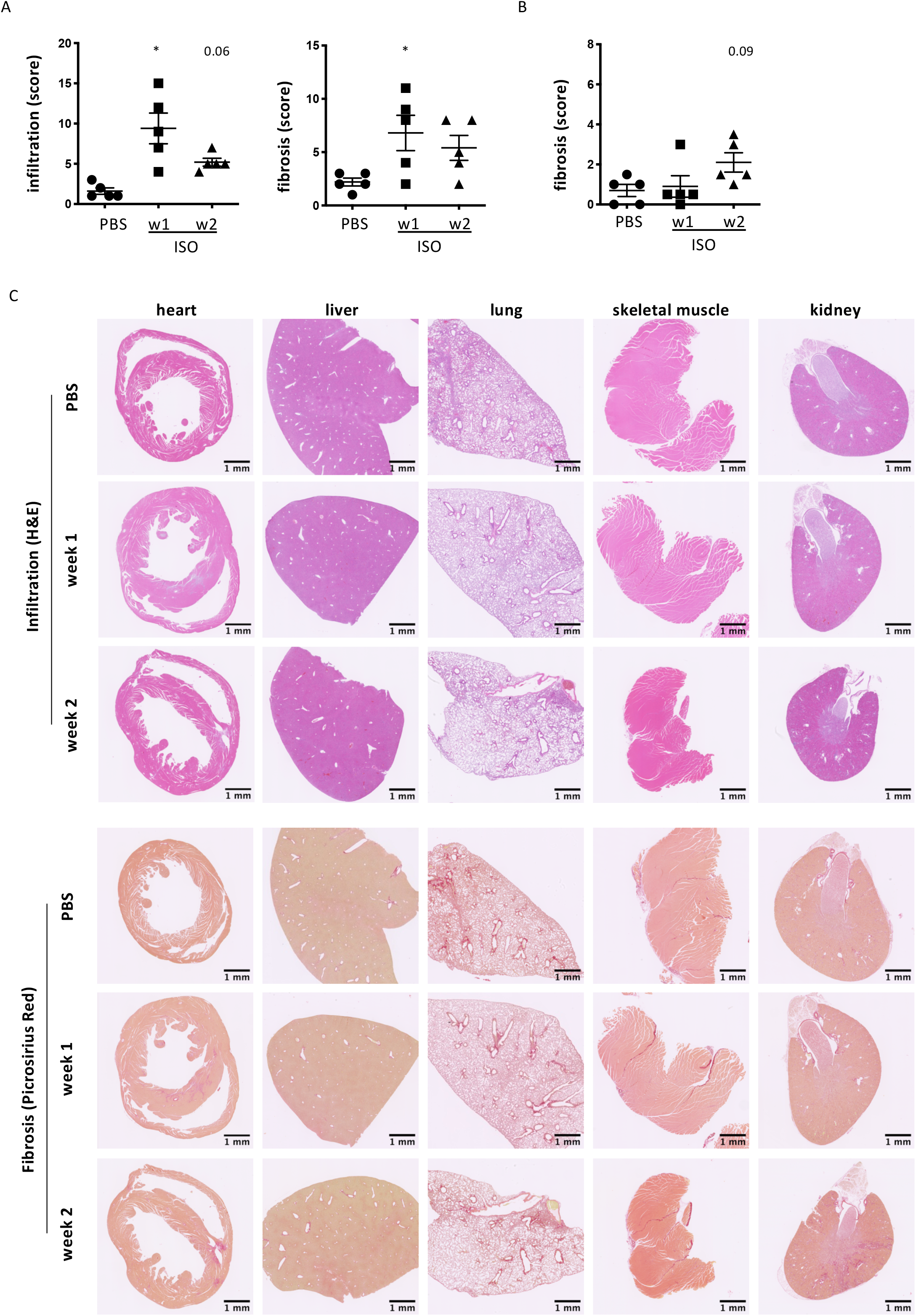
A single high dose of isoproterenol does not affect liver, lung and skeletal muscle, but pathological changes in the kidneys are observed secondary to cardiac damage at week 2. C57BL/6J mice were treated with a single one-off dose of isoproterenol to induce myocardial ischemia and cardiomyocyte necrosis. (A) Myocardial infiltration and fibrosis were scored on a scale from 0 to 3 (none, mild, moderate, severe) in 4 fields of view in 2 heart cross-sections at mid-level per mouse. (B) Kidney fibrosis was scored on a scale from 0 to 3 (none, mild, moderate, severe) in 4 fields of view in 2 cross-sections per mouse. n=5/group. Data are expressed as mean +/− s.e.m., *p<0.05 (non-parametric Kruskal-Wallis with Dunn's multiple comparisons post hoc test comparing each time point to baseline). (C) Micrographs of H&E- and Picrosirius Red-stained paraffin-embedded sections of post-isoproterenol organs for assessment of mononuclear cell infiltration (H&E) and interstitial fibrosis (Picrosirius Red). Representative image shown.

### (3) CD4+ T cells in heart-draining mediastinal lymph nodes are activated in response to a single dose of 160mg/kg isoproterenol

To investigate if the single high dose isoproterenol model was suitable for immunological readouts related to adaptive immune cell activation, we isolated heart-draining mediastinal lymph nodes for flow cytometry. 4 weeks after ISO injection, CD4+ helper T cells displayed an activated CD62L-CD44-effector cell phenotype (Figure 3).

### (4) A single dose of 160mg/kg isoproterenol triggers production of anti-heart auto-antibodies

Activated autoreactive CD4+ helper T cells are able to induce B cells to generate autoantibodies, which are a hallmark sign of an established autoimmune response. We therefore tested week 2 (acute) and week 12 (chronic) post-isoproterenol serum for the presence of anti-heart autoantibodies. We found a significant increase of auto-antibody levels at chronic stage after isoproterenol damage (Figure 4A). Importantly, these were of the mature class-switched IgG isotype indicating an established and persistent adaptive immune response against the heart. We also found a significant amount of IgG as well as immature IgM deposited in the myocardium (Figure 4B, C) indicating *in vivo* deposition. The deposition pattern followed the myocardial fibres and the inside of blood vessels indicating the potential for endothelial and cardiomyocyte damage.

**Figure 4:**
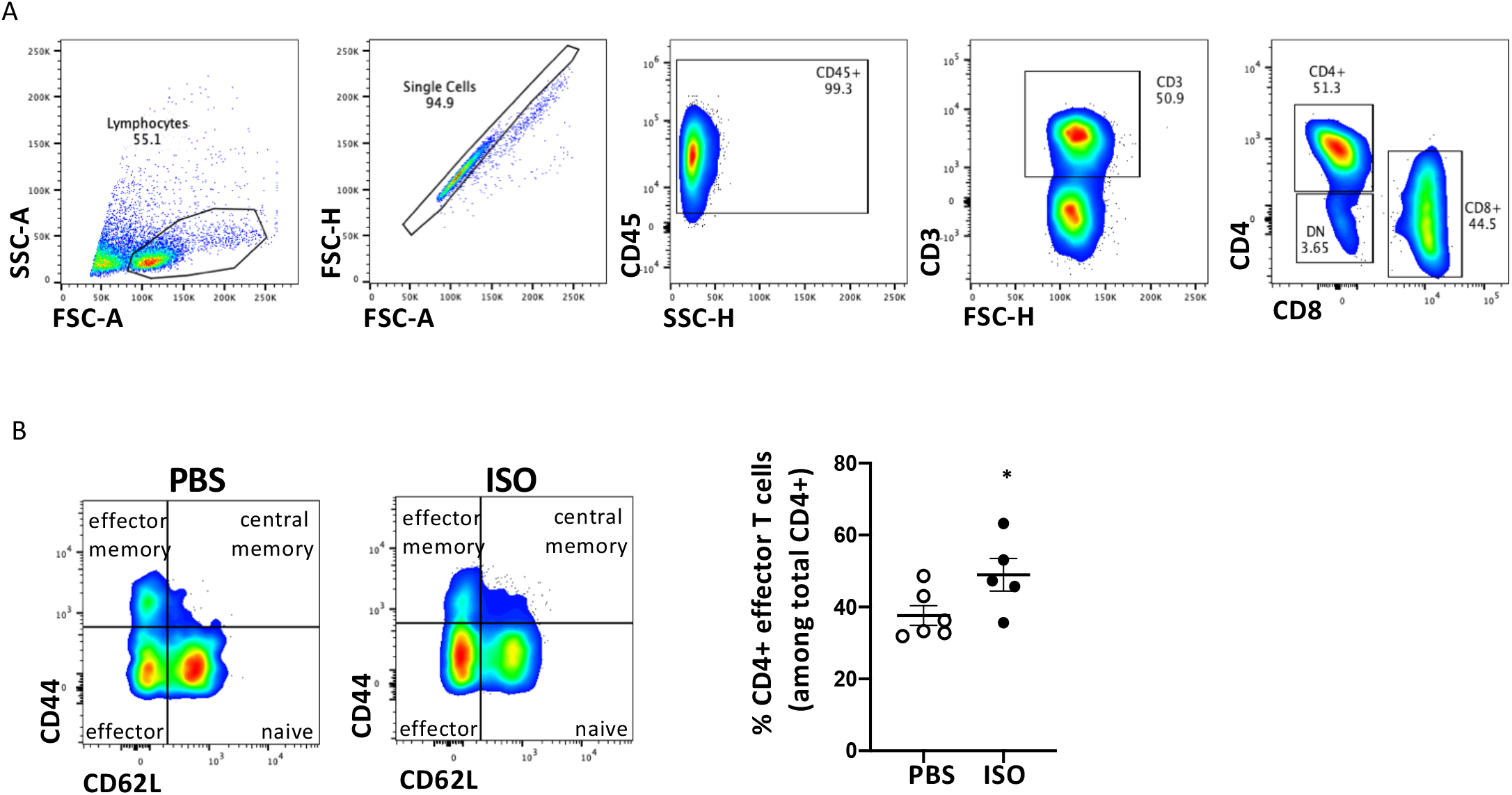
CD4+ T cells in heart-draining mediastinal lymph nodes are activated in response to A single dose of 160mg/kg isoproterenol. C57BL/6J mice were treated with a single one-off dose of isoproterenol to induce myocardial ischemia and cardiomyocyte necrosis and flow cytometry was performed on a single cell preparation of the mediastinal lymph nodes. (A) Gating strategy to obtain the CD3+ T cell population among CD45+ leukocytes. (B) Representative contour plots of CD62L and CD44 staining of CD3+CD4+ T cells and corresponding quantification of the DCD62L^−^CD44^−^ subpopulation of effector T cells of Isoproterenol-treated versus control mice. n=5-6/group. Data are expressed as mean +/− s.e.m., *p<0.05 (Student’s t test).

**Figure 5:**
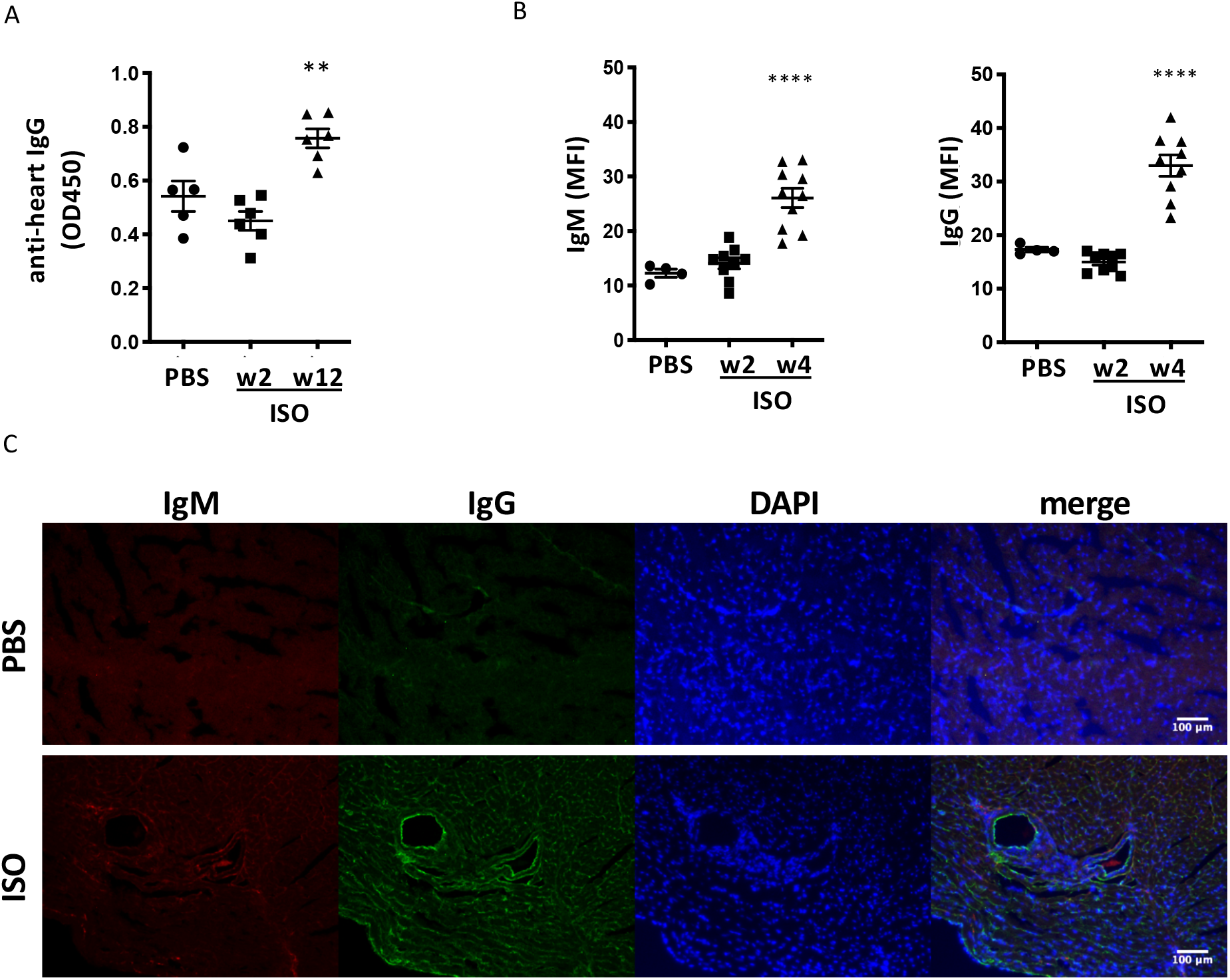
A single dose of 160mg/kg isoproterenol triggers production of anti-heart auto-antibodies. C57BL/6J mice were treated with a single one-off dose of 160mg/kg isoproterenol to induce myocardial ischemia and cardiomyocyte necrosis and the heart and serum were obtained for analysis of anti-heart autoantibody production. (A) Levels of anti-cardiac auto-antibodies in the serum over time as tested by ELISA using serum of ISOPROTERENOL-treated mice against rat cardiac lysate. (B, C) Quantification of MFI (B) and representative Immunofluorescence staining (C) of frozen heart sections of isoproterenol-treated mice 4 weeks after challenge, using goat anti-mouse IgG-FITC and rat anti-mouse IgM-PE to detect *in vivo* deposits of anti-heart auto-antibodies, green: IgG, red: IgM, blue: DAPI staining nuclei. n=5-6/group. Data are expressed as mean +/− s.e.m., **p<0.001 (one-way ANOVA with Dunnett's multiple comparisons post hoc test comparing each time point to baseline).

## Discussion

The definition of myocardial infarction differentiates patients with MI due to plaque rupture (T1MI) from those with myocardial necrosis due to oxygen supply-demand imbalance secondary to other acute illnesses (T2MI). Myocardial necrosis without symptoms or signs of myocardial ischaemia is classified as acute or chronic myocardial injury. Both myocardial injury and T2MI are common, yet these patients have poor short-term and long-term outcomes ^35^ with heart failure risk comparable to T1MI ^36^. Activation of the adaptive immune system and persistent autoimmunity against the heart have recently been suggested as pathological factors contributing to heart failure in response to MI ^37 14^. The number of corresponding experimental studies has surged and is anticipated to increase substantially in the near future as the cardiovascular community attempts to characterise the striking complexities of the post-MI immune response.

Here we show that a single dose of isoproterenol induces necrotic lesions in the heart. The initial trigger of necrotic cell death is most comparable to catecholamine-induced damage as seen during stress-related cardiomyopathies including takotsubo syndrome ^38 39^ as well as in severely-ill patients receiving catecholamines for cardiac pacing or inotropic support ^40^. High dose catecholamines also induce an imbalance between myocardial oxygen supply and demand, and the degree and pattern of inflammatory damage and replacement fibrosis is reminiscent of the myocardial damage incurred by a T2MI. Mild kidney fibrosis is also observed in isoproterenol-treated mice starting from week 2, which may be secondary to cardiac damage ^34^ as not apparent at earlier timepoints. Other organs are not affected, which confirms that off-target tissue damage is not a limiting factor in this model.

Importantly, isoproterenol-induced myocardial necrosis releases antigenic peptides which induce T cell activation in the heart-draining mediastinal lymph nodes. These activate B cells to produce mature IgG anti-heart autoantibodies, a sign of a bona fide autoimmune response against the heart. Anti-heart auto-antibodies indicating autoimmunity has thus far been demonstrated in a wide range of human heart conditions ^41^ and is likely due to high amounts of cardiac antigen in an inflammatory environment overwhelming immunological tolerance. Importantly, the adaptive immune system is activated by antigens released from necrotic cells irrespective of the initial cause of necrosis. Accordingly, we find anti-heart auto-antibodies in experimental models after surgical induction of MI via LAD ligation (Sintou et al. 2019) and under systemic inflammatory conditions ^28^, as well as in the present study after chemical induction of myocardial necrosis. The common factor in these situations is myocardial necrosis, which is sufficient to induce a downstream adaptive immune response against the heart.

Rodents are routinely used for experimental MI studies and infarcts are commonly induced by ligation of the left anterior descending (LAD) artery during open chest surgery, an invasive procedure associated with high mortality rates. Despite widespread uptake, this model is not free from limitations, which include additional tissue damage and inflammation triggered by invasive thoracotomy, an excessive severity of experimental infarcts and the use of young, healthy mice which do not reflect the average MI patient demographic well (Table 1). In particular, T2MI and other cases of more subtle ischaemic cardiac tissue injury are not reflected well by surgical LAD ligation. In addition, the isoproterenol model is particularly useful for readouts including chronic and peripheral immunological measures, e.g. serum levels of cytokines and anti-heart autoantibodies, phenotype and function of immune cells in peripheral lymphoid organs, as well as effects of therapeutic approaches on these aspects. Table 1 summarises characteristics of surgical MI and isoproterenol models in comparison to the clinical situation in human MI patients.

**Table 1:** Characteristics of LAD ligation and isoproterenol injection in comparison to human myocardial infarction.

|  |  |  |  |  |
| --- | --- | --- | --- | --- |
|  | Human myocardial infarction | Isoproterenol injection | Permanent LAD ligation | Temporary LAD ligation |
| Patient/subject population | Genetically diverse, various co-morbidities<sup>42</sup> | Mostly inbred, healthy, young animals<sup>43</sup><br>(experimental co-morbidities possible due to mild ISO challenge) | Mostly inbred, healthy, young animals<sup>43</sup><br>(experimental co-morbidities difficult as surgical challenge is already severe) |  |
| Cause of damage | Mixed population; temporary or permanent oxygen deprivation and/or I/R injury<sup>2</sup> | Overstimulation of β-adrenergic receptors inducing oxygen deprivation<sup>16 17</sup> | Permanent block of blood supply, surgical damage to thoracic cavity and pericardium<sup>44 45</sup> | Temporary block of blood supply - I/R injury, surgical damage to thoracic cavity and pericardium<sup>45 46</sup> |
| Early innate immune stimulus | tissue necrosis with/without complement activation<sup>47 48</sup> | tissue necrosis<sup>16 17 44</sup> |  | tissue necrosis and complement activation<sup>49</sup> |
| Severity | Varying severity, rarely transmural, mostly regional, sub-endocardial, micro-infarcts<sup>50 51</sup> | Diffuse patchy damage, primarily in endocardium<sup>16</sup> | severe trans-mural infarcts in LV (mimics severe T1MI without reperfusion ~25% of patients)<sup>52 53</sup> | Severe, often trans-mural infarcts in LV (mimics severe T1MI with reperfusion)<sup>54</sup> |
| Scarring | Varying degree of scarring depending on initial degree of damage<sup>55</sup> | Patches of fibrotic/scar tissue resembling T2MI or micro-infarcts<sup>16</sup> | majority of LV wall replaced by full thickness scar, additional scarring and damage due to MI surgery<sup>53 45</sup> |  |
| Functional and morphological response | Remodelling and functional decline to heart failure<sup>6</sup> | Mild effects on remodelling and dilation, mild functional decline<sup>30 56 57</sup> | strong remodelling response, dilation and progression to heart failure<sup>58 59</sup> | moderate remodelling response, dilation and progression to heart failure<sup>54 45</sup> |
| Adaptive immune stimulus | Cardiac antigen (e.g. myosin, troponin) in circulation<sup>60 61</sup> | Cardiac antigen (e.g. myosin, troponin) in circulation<sup>62 63</sup> |  |  |
| Adaptive immune response | Long-term auto-reactivity, anti-heart auto-antibodies<sup>64</sup> | anti-heart auto-antibodies<sup>14</sup> |  | Anticipated: anti-heart auto-antibodies |

Importantly, besides resource efficiency, replacing surgical MI procedures carries significant animal welfare benefits. Isoproterenol injections cause transient effects and are significantly less invasive than MI surgeries. In our hands, isoproterenol treatment carries an approximate mortality of 1% mostly due to acute arrhythmias in the first 24-48h after injection. Visible effects of appropriately titrated isoproterenol injections wear off within 1-2 hours after injection. Several opportunities to reduce necessary animal numbers also arise through the use of isoproterenol as alternative to LAD ligation surgery. Invasive thoracotomy causes a significant amount of tissue damage and inflammation and risks flawed interpretation of immunological readouts. Thus, there is scientific need for sham operated animals, while simple PBS injected controls are suitable for isoproterenol experiments. Isoproterenol is administered by intra-peritoneal injection, a simple technique that can be learned easily, avoiding increased mortality rates due to inexperienced researchers. Isoproterenol effects are also more homogenous between treated animals than surgical infarcts, allowing significant reduction of group sizes. Susceptibility to isoproterenol-induced damage varies between mouse strains. However, after appropriate dose studies, isoproterenol can be used in both sexes of all mouse strains, avoiding surplus animals and allowing the use of transgenic mice on their respective background.

In summary, a single high-dose isoproterenol injection is a suitable method to induce adaptive immune responses against the myocardium with significant additional resource and animal welfare benefits.

## Acknowledgements

We would like to thank staff at the animal facility at Imperial College London and The Jackson Laboratories for help with animal husbandry and maintenance. The authors also acknowledge the support by the Imperial College Facility for Imaging and Light Microscopy (FILM) and the LMS/NIHR Imperial Biomedical Research Centre Flow Cytometry. We gratefully acknowledge the contribution of Dr David Coleman and the Pathology Services Service at The Jackson Laboratory for expert assistance with the work described in this publication.

## Author contributions

E.F., M.G.H. and S.S. planned, designed, performed and analysed experiments. M.P., O.B. and J.Branca performed and analysed experiments. S.S. and F.S.N wrote the manuscript. J.Boyle and F.S.N. provided expert advice on histopathology and clinical cardiology and revised the manuscript. S.E.H., N.R. and S.S. provided supervision and financial support

